# Dietary and Developmental Shifts in Butterfly-Associated Bacterial Communities

**DOI:** 10.1101/196816

**Authors:** Kruttika Phalnikar, Krushnamegh Kunte, Deepa Agashe

## Abstract

Bacterial communities associated with insects can substantially influence host ecology, evolution and behavior. Host diet is a key factor that shapes bacterial communities, but the impact of dietary transitions across insect development is poorly understood. We analyzed bacterial communities of 12 butterfly species across different development stages, using 16S rDNA amplicon sequencing. Butterfly larvae typically consume leaves of a single host plant, whereas adults are more generalist nectar feeders. Thus, we expected bacterial communities to vary substantially across butterfly development. Surprisingly, very few species showed significant developmental transitions in bacterial communities, suggesting weak impacts of dietary transitions across butterfly development. On the other hand, bacterial communities were strongly influenced by butterfly species identity and dietary variation across species. Larvae of most butterfly species largely mirrored bacterial community composition of their diets, suggesting passive acquisition rather than active selection. Overall, our results suggest that although butterflies harbor distinct microbiomes across taxonomic groups and dietary guilds, the dramatic dietary shifts that occur during development do not impose strong selection to maintain distinct bacterial communities.

## INTRODUCTION

All animals are associated with bacterial communities, and this association can significantly affect host biology [1–4]. In particular, insect models have been instrumental in understanding the mechanisms by which bacterial partners influence host physiology [5–7]. Bacteria can provide various benefits to insects including efficient digestion, nutrient supplementation and detoxification; and can thus help insects to survive on suboptimal diets and occupy diverse dietary niches [7–13]. Through their impact on nutrient acquisition, bacteria can significantly alter traits such as host fecundity, survival and longevity [9,14–17]. In some insects, bacteria also influence other aspects of host physiology such as development, regulation of the immune system, hormone signaling and resistance against infections [7,15,18–20]. Furthermore, bacteria can affect the behavioral ecology of insects by influencing mate choice and social aggregation via pheromone production [21, 22]. Thus, to better understand insect ecology, evolution and behavior, it is important to determine the factors that affect insect-bacterial associations.

Diet is one of the key factors that shape bacterial community structure (diversity and abundance) across insect taxa [23–25]. For instance, gut microbiota of the omnivorous cockroach *Blattella germanica* is strongly affected by dietary protein content [26], and bacterial communities of *Drosophila melanogaster* vary with yeast, sugar and ethanol content in their diet [27]. A few studies also suggest that diet similarity is associated with convergent evolution of gut bacterial communities. For example, bacterial communities of detritivorous termites are more similar to detritivorous beetles and flies, rather than closely related wood-eating termites [23]. In contrast, in some cases diet does not significantly influence bacterial communities. For instance, communities of carnivorous and herbivorous larvae of different butterfly species are similar [28], as are communities associated with the adults and larvae of Emerald ash borers that feed on foliage vs. cambium [23]. Overall, diet has a variable but typically significant impact on bacterial communities of insects.

Dietary shifts occur not only across different insect taxa, but also within the lifetime of many insects including butterflies and moths (Lepidoptera), flies and mosquitoes (Diptera) and parasitoid wasps (Hymenopterans). In these orders, juvenile stages usually consume a distinct resource (typically solid) compared to the adult diet (typically liquid). These within-lifetime dietary switches are often associated with major developmental transformations. How do developmental transitions affect bacterial communities in these insects? Previous studies have found variable impacts of developmental stage on gut bacterial communities of insects [29–35]. However, there is no single comprehensive analysis of bacterial communities of an insect group. Important factors to investigate include multiple host developmental stages, individual variation between hosts, multiple host species to evaluate whether the results are generalizable, analysis of host diet to examine the route of bacterial acquisition, and analyses of wild-caught insects. Here, we attempt to fill these gaps using butterflies as a model.

Butterflies undergo complete metamorphosis involving four distinct developmental stages - egg, larva, pupa and adult. The dietary switch across stages is very stark in butterflies because larvae feed strictly on a solid diet, and adults only feed on liquids. The intermediate, non-feeding pupal stage is associated with massive tissue restructuring and physiological changes involving apoptosis and autophagy [36, 37]. During this stage, the larval gut (presumably a major reservoir of bacteria) is degraded and the adult gut is formed anew [38, 39]. Shortly after the adult butterfly ecloses from the pupa, it excretes metabolic waste, called meconium, generated in the pupal stage. Thus, within a butterfly’s lifespan, both diet and physiology undergo a major transition. In the butterfly *Heliconius erato*, bacterial communities vary significantly across development, with only few bacterial phylotypes shared between larvae and adults [40]. However, it is unclear whether similar patterns occur across butterfly species, given the immense diversity in their diet, behavior and life history.

We used 16S rDNA amplicon sequencing to characterize the bacterial community structure (composition and abundance of community members) of 12 butterfly species from 5 of the 6 described butterfly families: *Ariadne merione, Danaus chrysippus, Elymnias caudata, Spalgis epeus, Pseudozizeeria maha, Jamides celeno, Leptotes plinius, Gangara thyrsis, Erionata torus, Eurema blanda, Pieris brassicae* and *Papilio polytes*. Of these, we were able to analyze multiple individuals from different developmental stages of 9 species (Table 1). We also analyzed bacterial communities associated with the larval diets of 5 of these species to test whether larval bacterial communities are largely diet-derived (Table 1). We expected to find significant developmental changes in the bacterial community structure of each butterfly species, given the large differences in larval vs. adult diet of each host (Table 1). We also expected to find significant species-specific variation in bacterial communities associated with larvae, since each species has a distinct larval diet. On the other hand, we predicted that bacterial communities across adults would be more similar since they are thought to be generalists – multiple butterfly species can feed on similar resources (e.g. nectar from the same plant) while foraging.

**Table 1:**
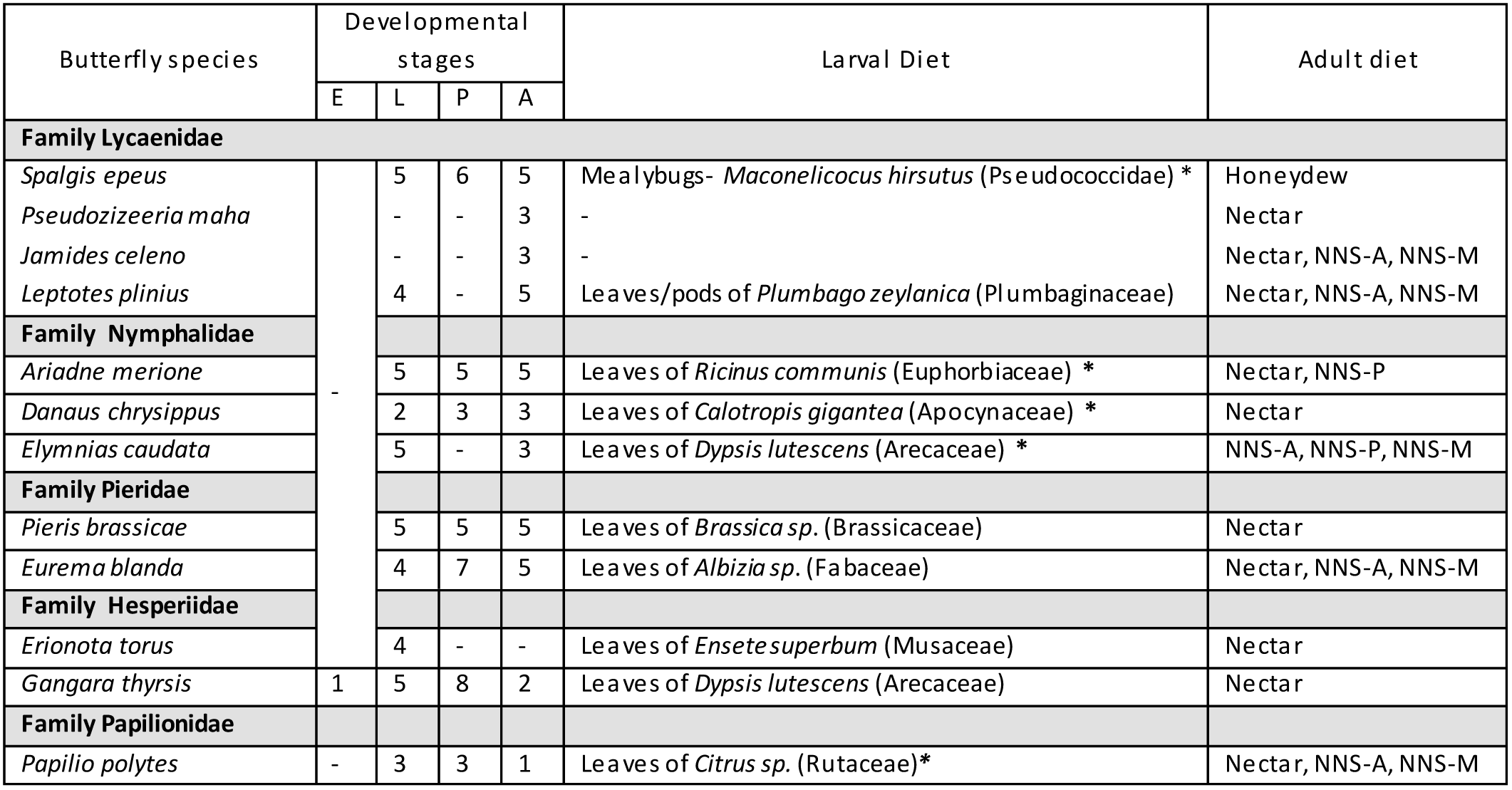
Butterfly sampling design. The table shows the number of replicates analyzed per developmental stage of each butterfly host species (E: Egg, L: Larva, P: Pupa, A: Adult). Asterisks indicate cases where we also analyzed bacterial communities associated with the larval diet. NNS-A, NNS-P and NNS-M correspond to **N**on **N**ectar **S**ources (NNS) used by butterfly adults. NNS-M: Adults obtain nutrients from mud fluids via **M**ud puddling. In most cases, males tend to mud puddle and not females. Thus, mud puddling may not be a prominent dietary resource for most adults in these samples as most are females. NNS-P: Adults feed on **P**lant products such as plant sap or rotting plant tissues (e.g. flowers and fruits). NNS-A: Adults feed on **A**nimal derived resources such as scat fluids, urine and sweat. All samples were collected from the same geographical location (NCBS campus and surrounding area, Bangalore) except samples of *P. brassicae* (Eaglenest wildlife sanctuary, Arunachal Pradesh).

Our work provides the first comprehensive analysis of bacterial communities across developmental stages of multiple wild-caught butterflies from different families and varied dietary habits. Unexpectedly, we find that within-species developmental dietary shifts have a weak impact on bacterial communities of butterflies. However, different host species harbor distinct bacterial communities, associated with dietary variation across hosts. Contrary to expectation, bacterial communities of generalist adult butterflies show slightly greater host-specificity compared to the communities of relatively specialist larvae. Together, our results indicate a weak impact of within-species dietary shifts but a stronger role of across-species diet divergence in shaping butterfly-associated bacterial communities.

## MATERIALS AND METHODS

### Sample collection and sequencing

We collected butterfly samples and larval dietary resources in and around the campus of the National Centre for Biological Sciences, Bangalore (NCBS) (13.0714° N,77.5802° E) from June 2014-January 2016, and from the Eagle Nest Wildlife sanctuary, Arunachal Pradesh (27°06′0″N 92°24′0″E) in June 2015. Information on butterfly identification, diets and larval host plants was taken from the Butterflies of India website (http://www.ifoundbutterflies.org/). We stored insects in 70% ethanol at −20°C until DNA extraction. We placed larvae and pupae in ethanol, and clipped off adult wings before storing the body. We surface-sterilized insect bodies by rinsing with fresh 70% ethanol and then 10% bleach for 30 seconds, each followed by three washes with sterile distilled water. We ground each sample in liquid nitrogen with single-use, sterilized pestles to homogenize the tissue. From the homogenized tissue, we e xtracted DNA using the Wizard® Genomic DNA Purification Kit (Promega) according to the manufacturer’s protocol. We dissolved the extracted DNA in sterile, DNAse/RNase free water and quantified DNA concentration using Nano-Drop (Nano-drop 2000, Thermo Fisher Scientific Inc., Wilmington, USA). We outsourced library preparation and Illumina sequencing to Genotypic Technology Pvt Ltd, Bangalore, India. We sequenced the V3-V4 hypervariable region of the 16S rRNA gene using 300 or 250 bp paired-end sequencing on the Illumina MiSeq platform.

### Data processing and analysis

We analyzed de-multiplexed MiSeq data using QIIME (version 1.9.1) [41]. We quality filtered reads with minimum quality score of q20. We removed chimeric sequences using USEARCH (version 6.1) [42] and assembled filtered reads into Operational Taxonomic Units (OTUs) with 97% sequence similarity using the UCLUST in QIIME. We picked OTUs with the ‘open reference OTU picking’ method in QIIME. After clustering the reads into OTUs, one sequence from each OTU was used as a representative sequence. We mapped representative sequences to the Green Genes 16S ribosomal gene database (Greengenes Database Consortium, version gg_13_5) to assign taxonomy using default QIIME parameters. We removed OTUs categorized as mitochondria and chloroplast prior to analysis. After obtaining the number of reads for each OTU in each sample, we further implemented 4 different cut-offs to remove rare bacterial OTUs from our analysis that may not contribute significantly to the bacterial community or may have arisen due to PCR or sequencing errors (see SI methods for details). These cut-offs are as follows:

1. We selected only the 5 most abundant OTUs for each sample, henceforth “Top 5 OTUs cut-off”
2. We eliminated OTUs that had <20 reads from each sample, henceforth “20 read cut-off”
3. We eliminated OTUs that had <5% relative abundance in at least 1 sample in our dataset, henceforth “5% abundance cut-off”
4. We eliminated OTUs that contributed <0.005% of the total reads across all samples, henceforth “0.005% abundance cut-off”

Although we considered all four rare-OTU filtering cut-offs for eliminating rare OTUs, we primarily focused on the most conservative top 5 OTUs cut-off and 5% abundance cut-off for our main analysis and results. We selected these abundant bacterial OTUs in order to capture the dynamics of potentially impactful community members. Another reason to restrict our analysis to dominant bacterial OTUs (top 5 OTUs and 5% abundance cut-off) was the inconsistency observed across sequencing runs of Illumina. Out of 130 samples, 42 samples were sequenced in one run and 88 samples were sequenced in a second flow cell. We found substantial variation in read depths in samples that were processed in these two independent rounds of Illumina sequencing, with large differences in the total number of OTUs found in replicate samples of the same host species analyzed in different sequencing runs (Figure S1). We suspect that the difference in OTU richness is due to differential sequencing depths and does not represent biologically relevant variation. To overcome this technical problem, we focused primarily on dominant bacterial OTUs. For each sample, the top 5 OTUs constituted a large proportion of the total bacterial community (average 87%, ranging from 66% to 99%; Figure S2), and hence we assume this is a fair representation of the 5 community. Similarly, with the 5% abundance cut-off we obtained comparable numbers of OTUs per sample across the two Illumina runs (Figure S1).

We characterized the bacterial communities associated with 120 butterfly samples from different families and varied dietary habits, and 10 samples of larval dietary resources (Table 1). We collected multiple developmental stages for 9 butterfly species and a single developmental stage for 3 species; all adult butterflies were females except for 4 adult samples where for 3 samples the sex was unknown and for 1 sample the adult was a male. After quality filtering data in QIIME (described above), we obtained a total of 2.6 × 10^7^ reads and 70348 OTUs across all 130 samples, with an average of 2× 10^5^ reads and 1931 OTUs per sample. We define OTUs as clusters of reads that have at least 97% sequence identity, and discuss all our results in terms of OTUs rather than taxonomically identifiable units. After applying the 4 different rare-OTU filters (top 5 OTU cut-off, 5% abundance cut-off, 0.005% cut-off and 20 reads cut-off), the total number of OTUs reduced to 5-13, 98, 964 and 11,364 respectively. In the analysis presented here, we focus on the two main cutoffs, 5% abundance cut-off and top 5 OTU cut-off (mentioned in each section as applicable).

To identify the most abundant and frequent OTUs across butterfly host and diet samples, we used the OTU subset obtained by applying the 5% abundance cut-off, representing the major bacterial taxa in each sample. For this subset, we obtained an average of 1.3 × 10^5^ reads and 43 OTUs per butterfly sample (ranging from 14 to 67 OTUs; Figure S3). To characterize the most frequent bacterial OTUs across all butterfly samples, we calculated the proportion of samples that harbored each bacterial OTU, and categorized all OTUs that were present in ≥ 80% of the samples as “frequent OTUs”. Similarly, to determine the most “abundant OTUs”, we calculated the relative number of reads contributed by each OTU (i.e. relative abundance) in each sample, and then calculated the average relative abundance of a given OTU across all samples. In addition, we identified the 5 most abundant bacterial phyla, classes, orders and families by calculating the proportion of reads contributed by the respective taxon out of the total reads across all butterfly samples.

We used R for all statistical analyses [43]. To determine whether bacterial community composition changes across development and across larvae and their diets, we performed Permutational Multivariate ANOVA (PERMANOVA) using the Adonis function in the R package Vegan [44]. To test the impact of developmental stage and host species on bacterial community composition, we used Canonical Analysis of Principal Coordinates based on Discriminant Analysis (CAPdiscrim) and the function Ordiellipse in the R package BiodiversityR (SI methods). We used non-parametric Kruskal Wallis tests in R to analyze variation in the relative abundance of specific bacterial OTUs across different samples (SI methods).

## RESULTS

### Butterfly microbiomes are dominated by the genus *Wolbachia* and families Methylobacteriaceae and Enterobacteriaceae

Across all butterfly samples (developmental stages of 12 species), OTUs from Methylobacteriaceae and Enterobacteriaceae were among the most frequent as well as abundant OTUs (Figures 1A & 1B). Bacterial OTU12 and OTU14 (family Methylobacteriaceae), OTU63 (genus *Staphylococcus*), OTU49 (genus *Micrococcus*) and OTU9 (family Enterobacteriaceae) were the most frequent OTUs, found in >80% of butterfly samples regardless of host species and developmental stage (Figure 1A). However, these common OTUs were not always the most abundant OTUs across butterflies. The most abundant OTUs were OTU3 (*Prevotella copri*), OTU1 and OTU2 (genus *Wolbachia*), OTU12 (family Methylobacteriaeceae) and OTU5 (genus *Capnocytophaga*) (Figure 1B). Notably, *Wolbachia* were both abundant and frequent across butterfly hosts (Figures 1A & 1B). At the phylum level, Proteobacteria, Firmicutes and Bacteroidetes together represented ∼95% of the total reads (Figure 1C). At the class level, Alphaproteobacteria, Gammaproteobacteria and Bacilli were the dominant bacterial groups (Figure 1D), whereas at the order level Rickettsiales, Enterobacteriales and Lactobacillales dominated the bacterial community (Figure 1E). The families Rickettsiaceae, Enterobacteriaceae and Prevotellaceae were most abundant and contributed >70% of the total reads (Figure 1F). These patterns are broadly similar to the findings of previous analyses of insect-associated microbiomes [23, 24]. However, the abundance of a bacterial taxon was not always proportional to the diversity of OTUs representing it. For example, although Firmicutes (11.9%) and Bacteroidetes (10.5%) had similar abundance (Figure 1C), Firmicutes were represented by 23 distinct OTUs whereas Bacteroidetes were represented by only 7 OTUs. Next, we tested whether these patterns of bacterial occurrence and abundance showed significant developmental and host-associated variation.

**Figure 1:**
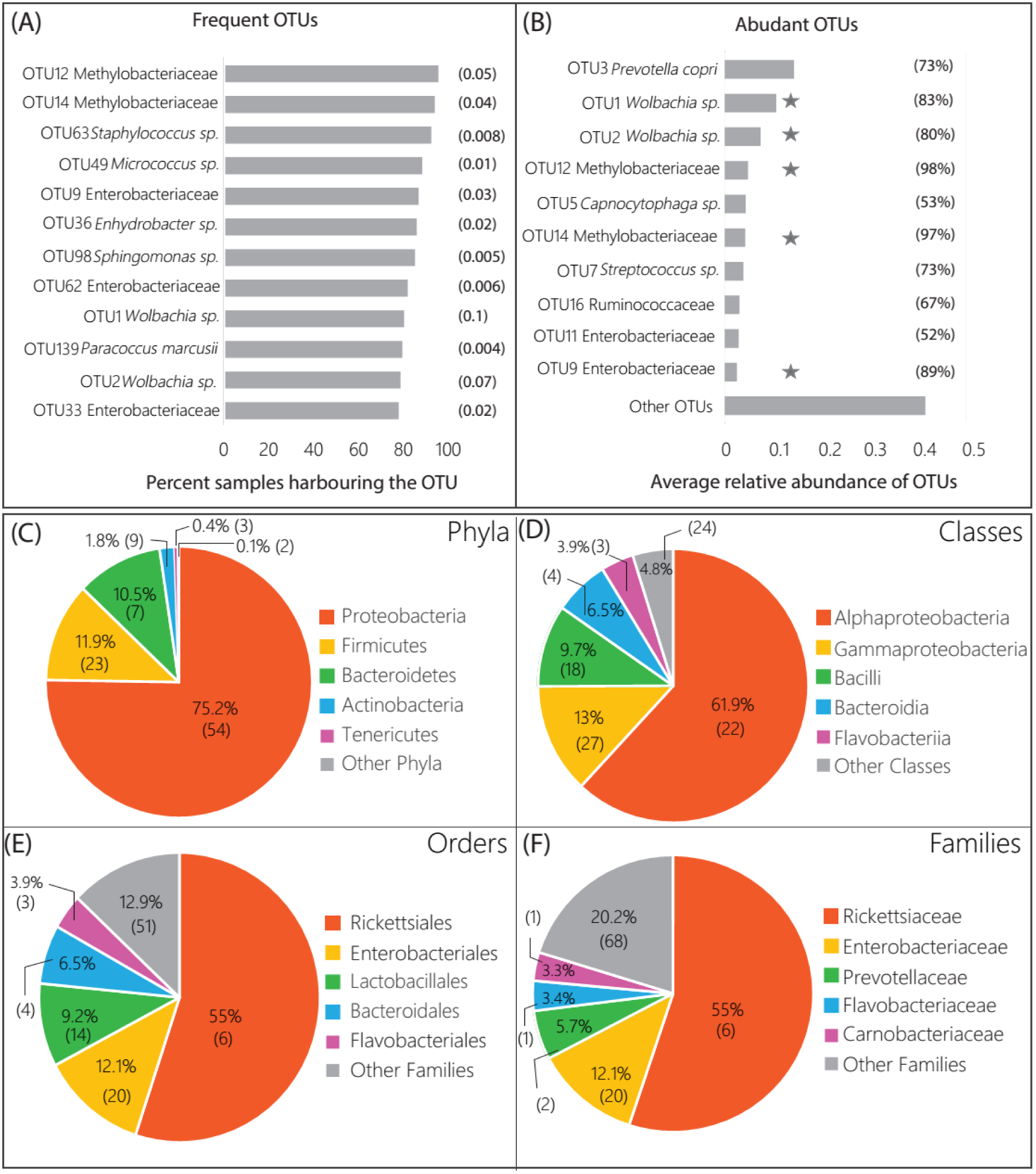
The most frequent and most abundant bacterial OTUs associated with butterflies. **(A)**The most frequent bacterial OTUs occurring in > 80% of butterfly samples. Numbers in parentheses represent the average relative abundance of each OTU. **(B)** The 10 most abundant bacterial OTUs (with the highest average relative abundance) across butterfly samples. Numbers in parentheses show the proportion of butterfly samples that harbored each abundant OTU. OTUs highlighted by a star are both abundant and frequent OTUs. **(C-F)** The five most abundant bacterial taxa across all butterfly samples. Each pie chart shows a different taxonomic rank, with each slice representing the percentage of total reads contributed by the taxon. Numbers in parentheses indicate the number of OTUs within each bacterial taxon

### Bacterial communities do not differ significantly across butterfly development

We evaluated the change in relative abundance of the top 5 bacterial OTUs across developmental stages of 9 butterfly species. Surprisingly, we found significant stage-specific variation in bacterial community composition and abundance only in 4 out of 9 butterfly species (Figure 2). When we tested the impact of developmental stage on bacterial community dynamics using alternative rare OTU cut-offs, we obtained similar results (Table S1). Note that we expected developmental variation in bacterial communities primarily due to the dietary shifts associated with development. Since the pupal stage is non-feeding, we next compared the bacterial communities of only larvae and adults of each species. In this analysis, only one butterfly species showed significant developmental variation in bacterial communities (*G. thyrsis*; PERMANOVA, p = 0.04, Figure 2D). An extreme example is the butterfly *L. plinius*, where the dominant bacterial community was almost identical across larvae and adults (Figure 2F). For most species, we did not find significant variation in diversity indices such as bacterial community richness and evenness across different life stages (Figure S4). Examining the dominant bacterial communities of replicate host samples, we found substantial individual variation in bacterial communities in nearly every developmental stage of each host species (Figure S5; Table S2). To test whether greater individual level variation may have masked the impact of development in some butterfly species, we calculated the coefficient of variation (CV) for each of the top 5 bacterial OTUs for each developmental stage of each host species (Figure S6). However, CV values were not significantly different across butterfly hosts that showed a significant vs. non-significant impact of developmental stage on bacterial communities. Hence, we cannot attribute the lack of a developmental signal on bacterial community composition to greater inter-individual variability in some hosts. Together, these results indicate very weak developmental variation in butterfly bacterial communities.

**Figure 2:**
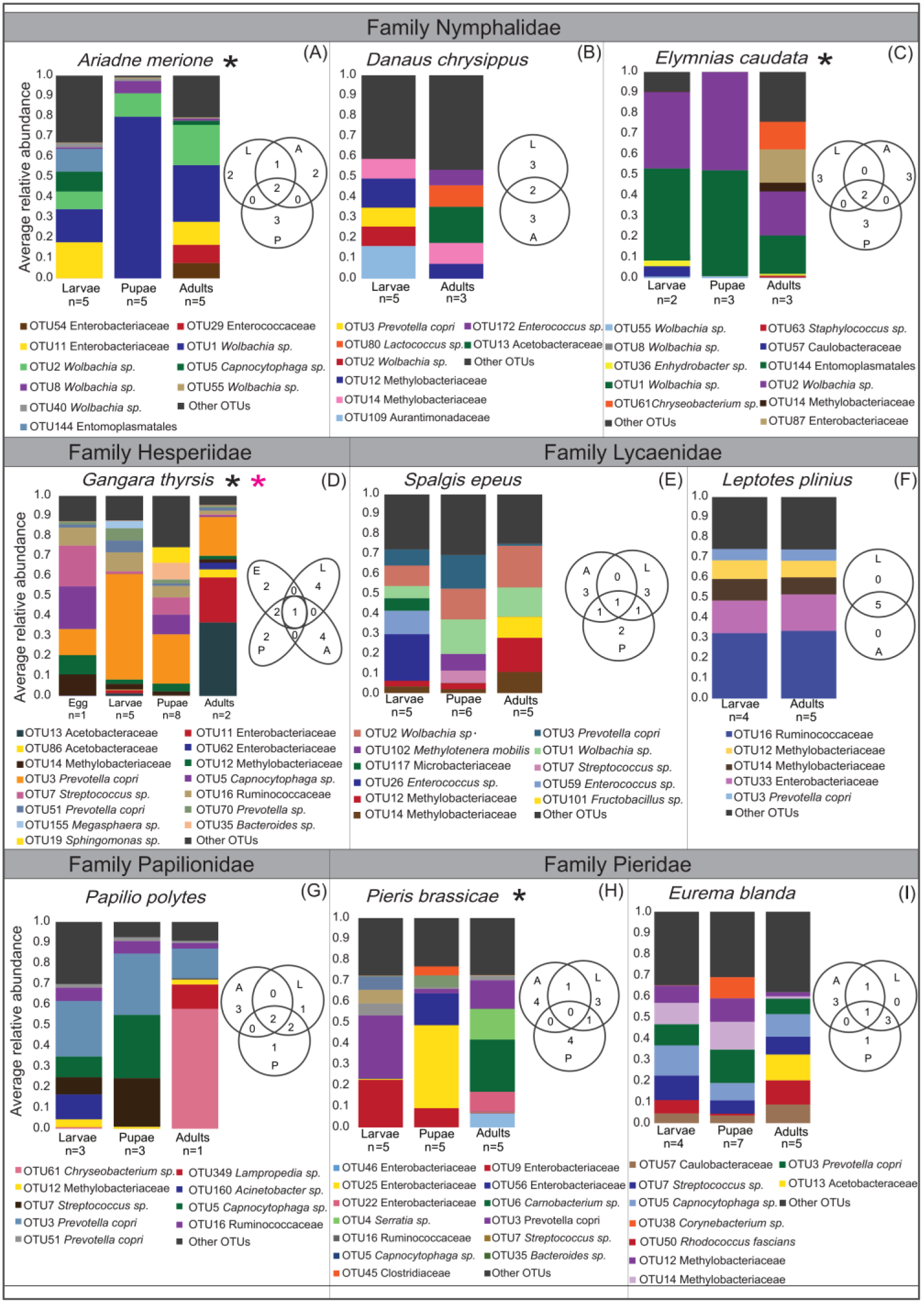
Variation in bacterial community composition across butterfly life stages. Stacked bar plots show the average relative abundance of the top 5 dominant bacterial OTUs across all developmental stages of a host. Each panel shows data for a single butterfly species; panels are grouped by family. Venn diagrams in each panel show the number of dominant OTUs that are shared or unique to each developmental stage. Asterisks (black) next to butterfly species names indicate significant variation in bacterial community composition across all developmental stages (Permutational Multivariate ANOVA PERMANOVA with 10000 permutations; p<0.05). Asterisks (pink) indicates significant variation in bacterial communities between larvae and adults

### Butterfly-associated bacterial communities are host-specific

Next, we used CAPdiscrim and PERMANOVA (see methods) to test the effect of host species and taxonomic family on the dominant bacterial communities of butterflies (5% most abundant OTUs). With CAPdiscrim, we obtained between-class variance, whereas with PERMANOVA we calculated the overall variance explained. To visualize the overlap in bacterial communities across host species and families, we used 95% confidence ellipses for each group. Within each developmental stage, we found significant variation in bacterial communities across butterfly species (MANOVA, p<0.05; Figure 3), with the first two linear discriminants (LD1+LD2) capturing >80% between-group variation (Table S3A). The total between-group variation explained (LD1+LD2) was highest for pupae (98%) and lower for larvae (88%) and adults (85%) (Table S3A). However, when using other rare OTU filters, we observed that total LD explained was greater for adults than larvae (Table S3A). Similarly, a PERMANOVA analysis showed that host species identity explained slightly more variation in the microbiomes of adults (∼8% more than larvae; Table S4). Finally, the impact of host species was weakened if we pooled individuals across developmental stages (Tables S3A and S4; Figure S7). Thus, contrary to expectation, we observed stronger host-specificity in bacterial communities associated with adult butterflies rather than larvae.

**Figure 3:**
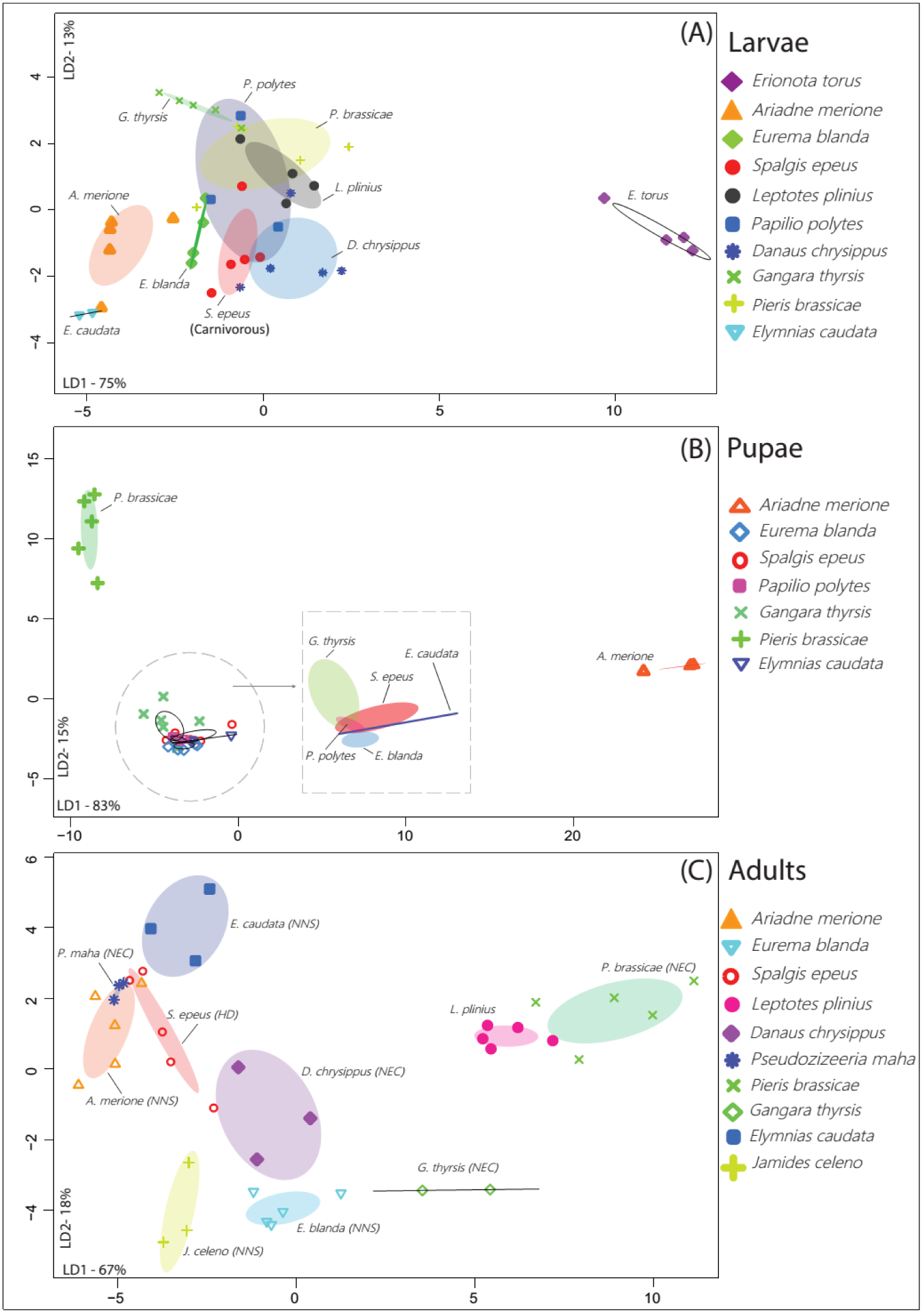
Variation in bacterial communities of developmental stages of different butterfly host species. Panels show Constrained Analysis of Principal Coordinates (CAP) of larvae (A), pupae (B) and adults (C) based on the composition and relative abundance of bacterial OTUs after applying a 5% abundance cut- off. In each panel, different colors and symbols indicate distinct butterfly species. Axis labels indicate the proportion of between-group variance (%) explained by the first two linear discriminants (LD1 and LD2). Ellipses represent 95% confidence intervals. For each panel, we observe a significant effect of host species (p<0.05, multivariate ANOVA). Carnivorous *S. epeus* larvae (Panel A) and the adult dietary resource (see Table 1) of each butterfly species are marked (Panel C). In panel A, the impact of host species remained significant even after removing the potential outlier *E. torus* larvae (Figure S12; MANOVA, p<0.05).

As with host species, host taxonomic family also significantly affected bacterial community structure (Tables S3A and S4; Figure 4). For larvae (Figure 4A), family Papilionidae showed substantial overlaps with other families based on 95% confidence ellipses. However, family Papilionidae was represented by only 1 species (*P. polytes*). When we performed the same analysis without family Papilionidae, we observed distinct clusters of larval families with some overlap between family Lycaeniade and Pieridae (Figure S8). Unlike the patterns observed with host species, the total variation explained by host family was not different for larvae and adults (Table S4). Comparing the relative impact of host species and family on community composition in each developmental stage, we found that host species typically explained greater total variation (∼30% more for larvae and adults, and ∼10% more for pupae; Table S4). This suggests that bacterial communities are more strongly structured by host species rather than host family. These patterns were generally consistent even if we removed the potential endosymbiont *Wolbachia* from our analysis (Figures S9, S10; Table S3B); although removing *Wolbachia* decreased the variation explained by total LD (Table S3B) and reduced the impact of host family on larval bacterial communities (MANOVA, p = 0.06).

**Figure 4:**
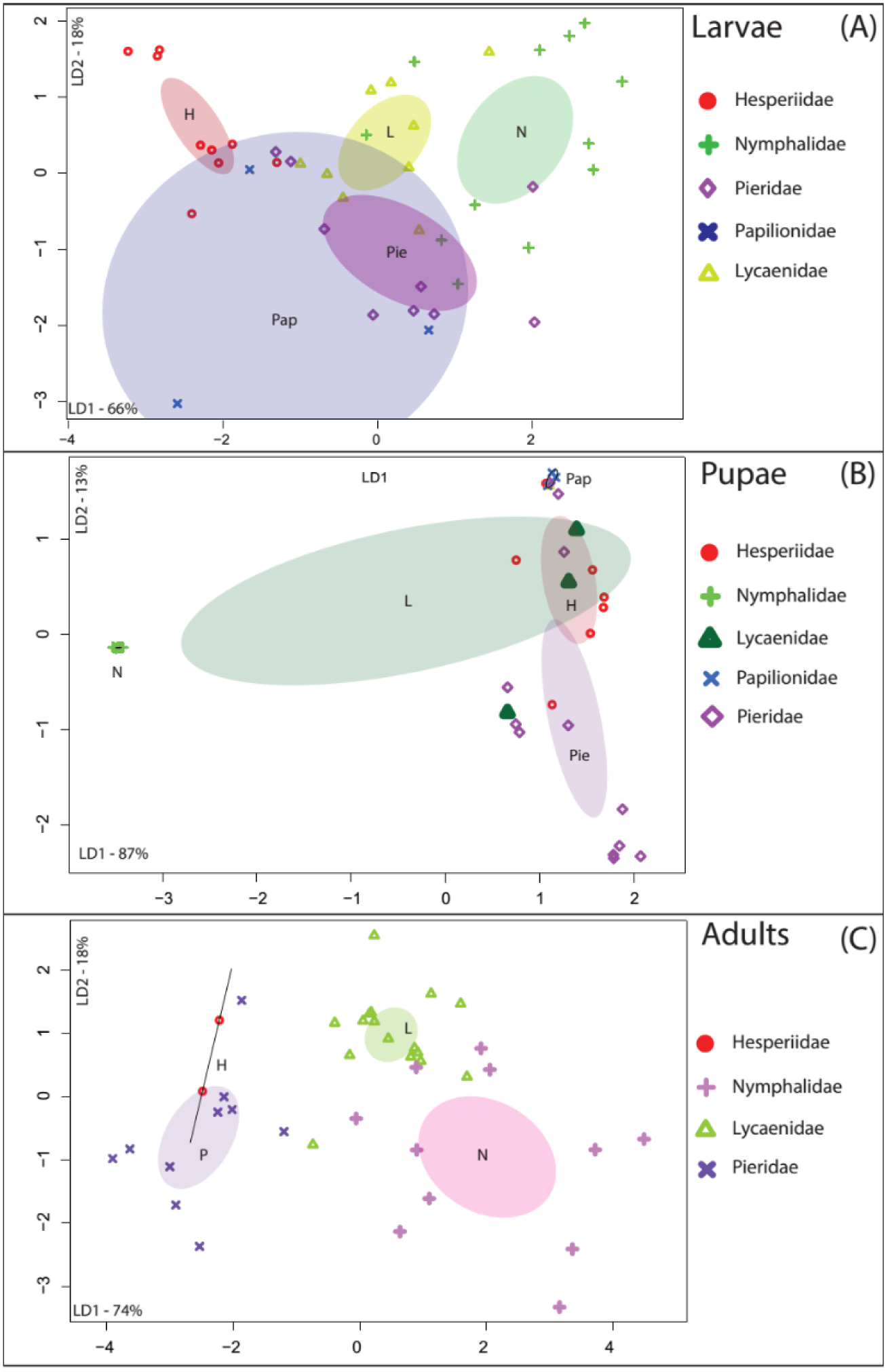
Variation in bacterial communities across butterfly host families. Panels show Constrained Analysis of Principal Coordinates (CAP) of larvae, pupae and adults based on the composition and relative abundance of bacterial OTUs after applying a 5% abundance cut-off. We pooled all individuals belonging to a butterfly taxonomic family, regardless of their species. In each panel, different colors and symbol indicate distinct butterfly families. Axis labels indicate the proportion of between-group variance (%) explained by the first two linear discriminants (LD1 and LD2). Ellipses represent 95% confidence intervals. For each panel, we observe a significant effect of host family (p<0.05, multivariate ANOVA)

### Diet affects bacterial communities across butterfly host species

The strong host-specificity observed in bacterial communities could arise through neutral mechanisms (e.g. distinct input communities due to dietary differences), via host-imposed selection (e.g. due to dietary or physiological differences), or a combination of both processes. To determine the importance of dietary differences (through neutral or selective mechanisms of community assembly), we compared bacterial communities associated with larval dietary sources (Table 1, Figure 5A). We found that the average Bray-Curtis dissimilarity in bacterial communities across larval dietary resources was ∼40% (Table S5). For dietary sources with more replicates (Table 1), a PERMANOVA analysis gave similar results (*Df =3, R*^2^ *= 0.73105, p = 0.0379*). These results indicate a potential role for larval diets in driving host-specific bacterial communities. Interestingly, in 5 of 6 species, ∼80% of the OTUs found in dietary resources were also found in the dominant bacterial communities of larvae (Figure 6, Table S6). Hence, it is not surprising that the bacterial community structure of larvae and their diets did not vary significantly (Figure 6; PERMANOVA, p>0.05) except in *G. thyrsis* (PERMANOVA, p=0.047). Notably, the carnivorous larvae of *S. epeus* shared ∼75% of their bacterial community with that of their insect prey *M. hirsutus* (mealybugs) and ∼50% of the community with the mealybugs’ host plant *Hibiscus* (Figure 6D; Table S6). Overall, our results suggest that the bacterial communities of butterfly larvae are largely shaped by their diet. Note that in contrast to a recent study of Lepidopteran caterpillars where >80% of the 16S reads were attributed to diet (plant)-derived chloroplast and mitochondria [57], we found that in our samples this proportion was only about 33% (Figure S11).

**Figure 5:**
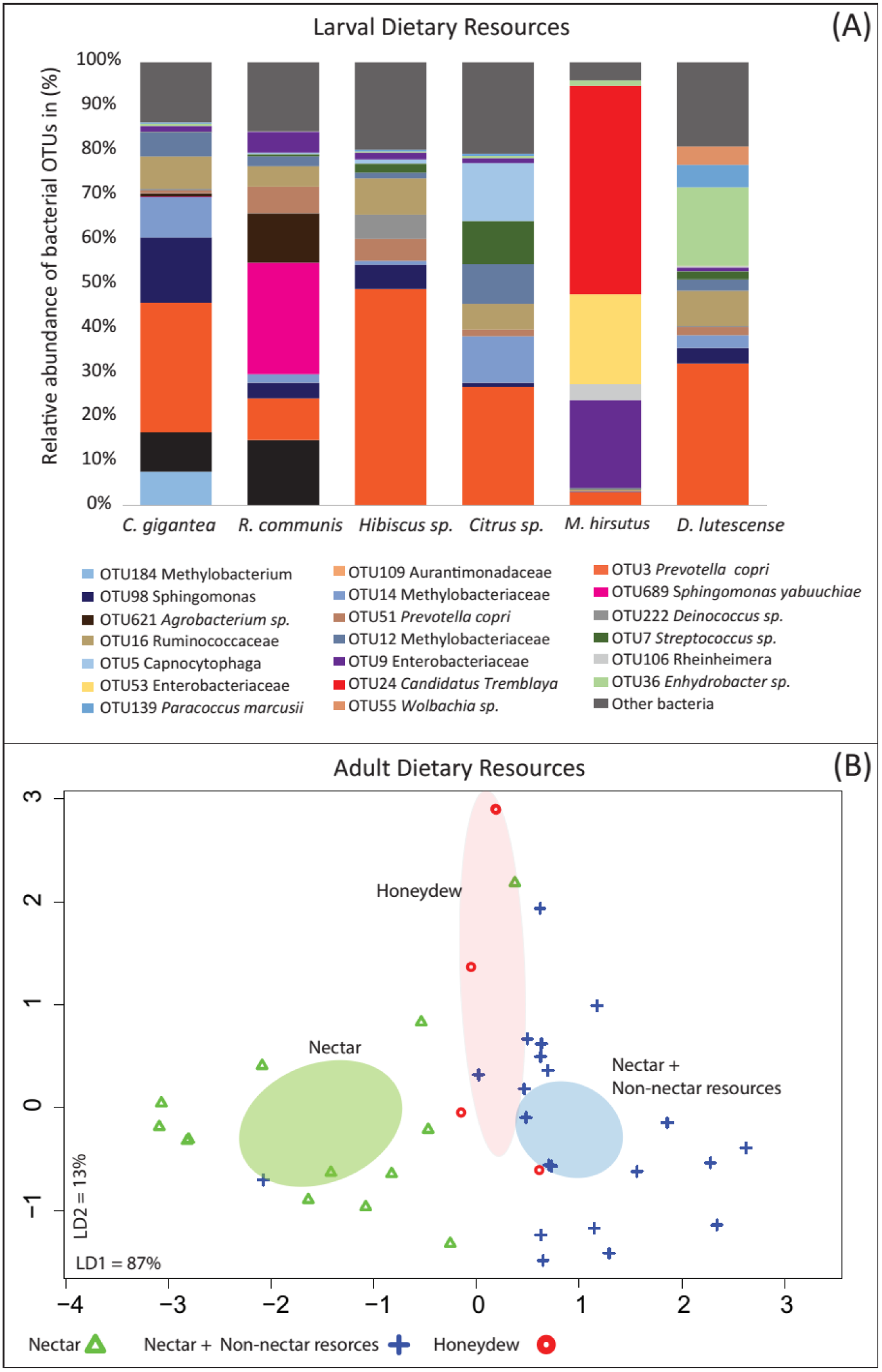
Variation in bacterial community composition across butterfly dietary resources. **Panel A**-Stacked bar plots show the average relative abundance of the top 5 dominant bacterial OTUs across all larval dietary resources. **Panel B-** Variation in bacterial communities of adults with different dietary habits (see Table 1). Panel shows Constrained Analysis of Principal Coordinates (CAP) of adults from different dietary guilds based on the composition and relative abundance of bacterial OTUs after applying a 5% abundance cut-off. Axis labels indicate the proportion of between-group variance (%) explained by the first two linear discriminants (LD1 and LD2). Ellipses represent 95% confidence intervals. We observe a significant effect of host family (p<0.05, multivariate ANOVA).

For wild-caught adults, we had no information on specific dietary resources. However, we compared bacterial communities across host species known to vary in their dietary habits (Table 1; Figure 5B). Our adult butterfly species included 3 broad dietary types, with species that feed on (a) nectar only, (b) nectar + non-nectar resources and (c) honeydew (Table 1). A CAPdiscrim analysis showed significant variation across these groups (LD1+LD2 = 100%, classification success = 79%, MANOVA p = 0.007; Figure 5B), as did a PERMANOVA analysis (*Df =2, R*^2^ *= 0.092, p= 0.0054*). In fact, we observed significant host specificity within the dietary guild with multiple host species (PERMANOVA for species with NNS diet in Table 1; *df = 4, R*^2^ *= 0.39, p=0.0002*), suggesting that each of these species may use distinct non-nectar resources. Note that we removed *Wolbachia* OTUs for this analysis, since they may be non-gut associated endosymbionts. Thus, adult dietary groups may strongly influence bacterial communities across butterfly species (Figure 5B). Together, our results suggest that dietary variation may explain the observed host-specificity in butterfly associated bacterial communities. However, whether the impact of diet reflects neutral (passive acquisition) or host-mediated selective processes remains unclear.

**Figure 6:**
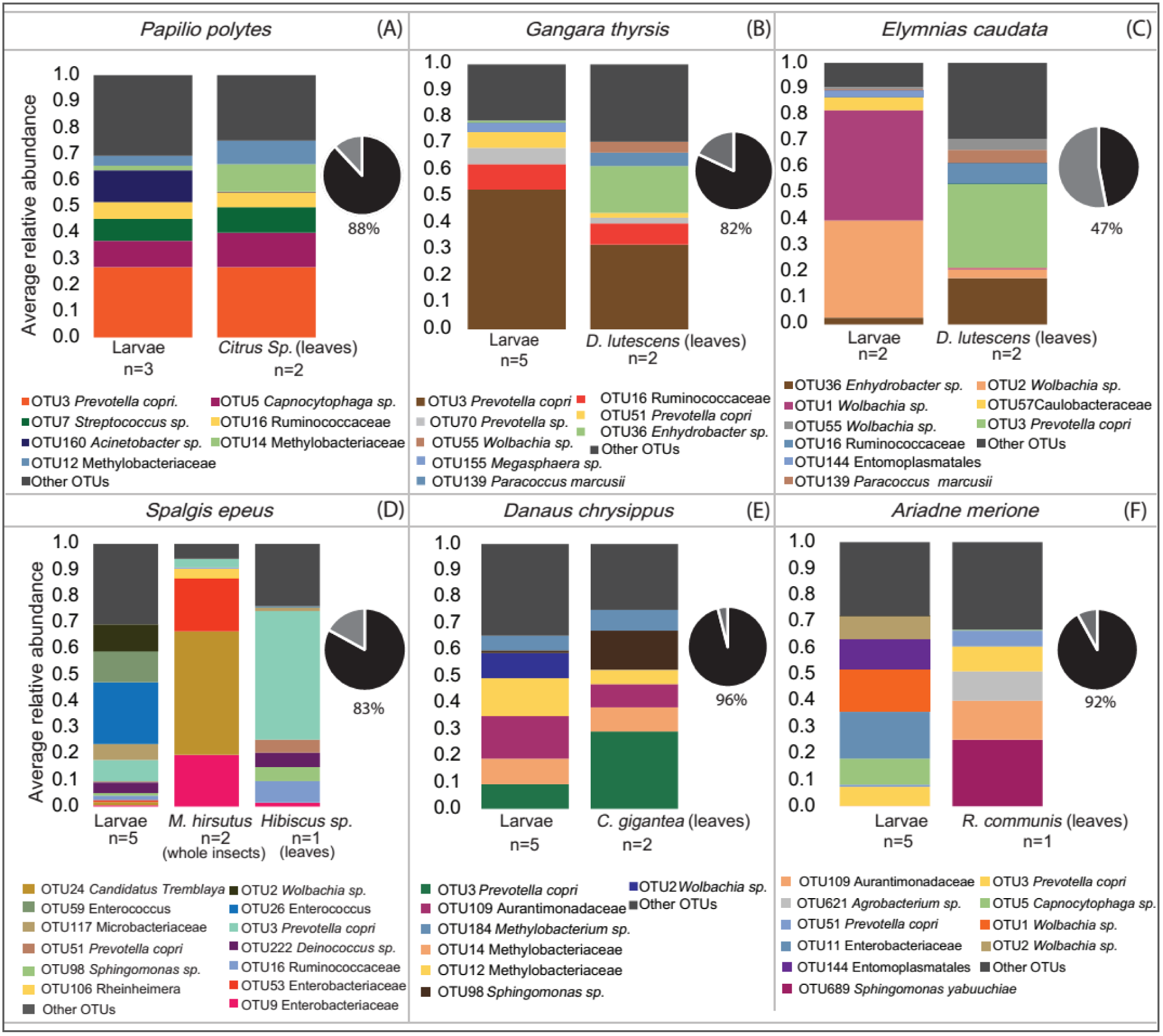
Bacterial community composition of larvae and their diets. Stacked bar plots represent the average relative abundance of the 5 most abundant bacterial OTUs. Each bar represents a larval stage or a dietary resource; n = number of replicates sampled. Each panel shows data for a different butterfly species. Variation in bacterial community composition was tested using permutational multivariate ANOVA (PERMANOVA, 1000 permutations). For *G. thyrsis*, p<0.05; for all other comparisons, p>0.05. Pie charts in each panel represent the proportion (%) of dietary bacteria (OTUs) found in larvae (black slice) at the 5% relative abundance cutoff.

## DISCUSSION

Across animals, a large body of work shows that diet plays a major role in shaping host-associated microbial communities [45–50]. We demonstrate that although this pattern holds across butterfly species, the dramatic dietary shifts across developmental stages of a given host have a very weak role in shaping bacterial community structure. This is surprising because differences in diet quality and physiological variation across metamorphosis should generate strong selection to maintain distinct sets of beneficial microbes in each life stage. For instance, the larval diet (mostly leaves) can be difficult to digest as it is typically enriched in cellulose, hemicellulose, lignin, pectin [51], several toxic plant defense compounds [52], and is often nitrogen limited [7]. Conversely, the adult diet (primarily nectar for species in this study) is typically composed of sucrose, glucose, hexose, fructose and amino acids [53, 54] with fewer or no toxins since plants attract pollinators with the nectar reward. In spite of this stark variation in diet, we did not find significant changes in bacterial communities across larvae and adults for most butterflies.

The overall lack of developmental signal could arise via two mechanisms. First, larvae and adults of a given host species could independently acquire similar bacteria from their respective diets. Second, a large proportion of larval bacterial community may be maintained across metamorphosis, causing significant overlap in bacterial communities of larvae and adults. The first scenario is less likely, because previous reports show that leaves and nectar of different plant species harbor distinct bacterial communities [59-62]; thus it is unlikely that larvae and adults would take up similar bacteria from their respective diets. However, we found some support for the second hypothesis. Analysis of pupal bacterial communities showed that in some butterflies (3 of 7 species with pupal sampling), pupae had similar bacterial composition and richness as that of the larvae and adults (Figure 2, Figure S3). This is in contrast to previous reports with other insects, where pupae seem to harbor fewer OTUs relative to larvae and adults [29,31,34,40,55]. In fact, in some cases the diversity and abundance of bacterial OTUs increased during the pupal stage (Figure 2, Figure S3). For instance, in *P. polytes*, the average relative abundance of OTU7 (*Streptococcus sp.*) increased roughly 200% from larvae to pupae, and in *P. brassicae* the average relative abundance of OTU25 and OTU56 (family Enterobacteriaceae) increased by 45% and 85% respectively from larvae to pupae. Similarly, in *S. epeus* and *G. thyrsis*, OTU richness in pupae was higher than in larvae and adults. An interesting avenue for further work is to identify where and how bacteria are maintained or enriched in pupae, and whether the enrichment is beneficial for the host. This is especially relevant for understanding the rare instances where we observed significant changes in bacterial communities across developmental stages.

Unlike developmental stage, we found that host species and taxonomic family strongly impacted bacterial community structure. An earlier analysis of bacterial communities across 39 insect species from 28 families found similar results, showing that closely related insect taxa have more similar bacterial communities [71]. For butterflies, a strong impact of host species was expected because diet varies substantially across host species. However, we anticipated that larvae of different species would be much more distinct from each other than adults. This is because bacterial communities vary significantly across plant species [58, 59, 60], and should be reflected in distinct larval microbiomes even in the absence of selection. Specifically, larvae of our focal butterfly species feed on distinct host plants with distinct microbiomes. On the other hand, adults of most butterfly species are generalists [56, 57]. Therefore, we expected specific associations between larvae and bacterial partners but weaker associations for adults. To the contrary, we found that the impact of host species on bacterial communities of larvae and adults was slightly greater in adults. For larvae, we found substantial overlap with the bacterial communities associated with larval diet, indicating that larval bacterial communities are largely shaped by passive acquisition of bacteria from dietary resources. This pattern is consistent with weak selection for host-bacterial associations in butterfly larvae, as suggested by a recent analysis of Lepidopteran (mostly moth) larvae [58]. The stronger signal of host-specificity in adults could arise if different species specialize on different nectar or non-nectar resources, acquiring distinct sets of bacteria [61, 62]. Indeed, we observed that broad dietary groups explained a large proportion of variation in bacterial community structure in adults. In addition to species identity, taxonomic family also significantly impacted bacterial community structure across butterflies. This impact can arise due to several factors such as divergent, family-specific host physiology and immune system, potentially selecting or enriching for distinct sets of bacteria. Further work is necessary to test these hypotheses and determine the relative role of neutral and selective processes in shaping butterfly-associated bacterial communities.

At the level of bacterial taxonomy, our findings are generally congruent with previous work on insect-associated bacterial communities. The most abundant and frequent OTUs in butterflies are also common members of other insect-associated microbiomes. For instance, of the 12 frequent bacterial OTUs found across butterflies, families Methylobacteriaceae, Enterobacteriaceae and genus *Wolbachia* together represented 7 OTUs. These bacteria also dominate bacterial communities associated with insects from multiple orders [24], including Lycaenid butterfly larvae [28]. Similarly, the phyla Proteobacteria, Firmicutes, Bacteroidetes, and Actinobacteria accounted for ∼94% of the total bacterial abundance in butterflies; these phyla are again commonly observed in multiple insects [23, 24, 71] including the butterfly *Heliconius erato* [40] and butterfly larvae from family Lycaenidae [28]. Such widespread insect-bacterial co-occurrence may represent evolved functional relationships, allowing the bacteria to easily colonize and proliferate within a wide range of insects including butterflies. On the other hand, they may simply reflect the fact that these bacteria are commonly found in the phyllosphere and soil [72, 73] that serve as ecological or dietary niches for many insects including butterflies. Further work is necessary to distinguish between these hypotheses.

A large body of work has analyzed microbial communities associated with insects [18,23,24,74–76], but very few studies have investigated butterflies. There are about 19,000 species of butterflies worldwide [77] that exhibit large diversity in their dietary habits, ecological niches and life history. They are important pollinators and herbivores, while some are pests in the larval stages [78–80]. Butterflies are also used as model systems across several disciplines of biology such as genetics, behavioral ecology, developmental biology and evolutionary biology. Despite being such an ecologically important insect group and a widely used model system, very little is known about the bacterial communities associated with butterflies. Our study is one of the first investigations of bacterial communities harbored by a diverse set of wild-caught butterfly species, across developmental stages and larval diets. Surprisingly, we find that despite the large dietary switch, butterfly developmental stages have fairly similar bacterial communities, with weak evidence of specific co-evolved associations. Our work highlights the importance of comparative analyses across multiple species within an insect group, because focusing on any one butterfly species might have led to different conclusions. Further studies can build upon our results by including butterfly species with more diverse diets and better resolution across developmental stages; and by conducting manipulative experiments to test whether butterfly-bacterial associations are mutualistic, commensal or parasitic.

## ACKNOWLEDGEMENTS

This work was supported by funding from the National Centre for Biological Sciences (NCBS), ICGEB (grant CRP/IND14-01), the University Grants Commission India (Research fellowship to KP), and the Department of Science and Technology, India (Inspire Faculty Award IFA13 LSBM-64 to DA). We thank Rittik Deb, Aparna Agarwal, Gaurav Diwan and Umesh Mohan for help with QIIME and R analysis and for important suggestions. We thank members of the Kunte lab for help with butterfly identification and fieldwork. We thank Riddhi Deshmukh, Sruthi Krishnamurthy and Tashi for help with sample collection. For sample collection in the Eaglenest Wildlife sanctuary, research and collection permits were issued by the state forest department of Arunachal Pradesh, India (permit no. CWL/G/13(95)/2011-12/Pt-III/2466-70, dated 16/02/2015). We thank all staff at the field station at the Eaglenest wildlife sanctuary for their help during sample collection.

## DATA ARCHIVING

Raw Fastq files generated through illumine MiSeq amplicon sequencing will be uploaded in the European Nucleotide Archive (EMBL EBI) database (http://www.ebi.ac.uk/) under accession number PRJEB21255.

## AUTHOR CONTRIBUTIONS

DA and KP conceived of the project; KP, DA and KK designed and carried out the work; KP and DA analyzed data and wrote the manuscript with input from KK. All authors approved the final version of the manuscript.

